# Prediction of mammalian tissue-specific CLOCK-BMAL1 binding to E-box motifs

**DOI:** 10.1101/2022.06.27.497767

**Authors:** Daniel Marri, David Filipovic, Omar Kana, Shelley Tischkau, Sudin Bhattacharya

## Abstract

The mammalian circadian clock is based on a core intracellular gene regulatory network, coordinated by communication between the central nervous system and peripheral tissues like the liver. Transcriptional and translational feedback loops underlie the molecular mechanism of circadian oscillation and generate its 24 h periodicity. The Brain and muscle Arnt-like protein-1 (Bmal1) forms a heterodimer with Circadian Locomotor Output Cycles Kaput (Clock) that binds to E-box gene regulatory elements, activating transcription of clock genes. In this work we aimed to develop a predictive model of genome-wide CLOCK-BMAL1 binding to E-box motifs. We found over-representation of the canonical E-box motif **CACGTG** in BMAL1-bound regions in accessible chromatin of the mouse liver, heart and kidney. We developed three different tissue-specific machine learning models based on DNA sequence, DNA sequence plus DNA shape, and DNA sequence and shape plus histone modifications. Combining DNA sequence with DNA shape and histone modification features yielded improved transcription factor binding site prediction. Further, we identified the genomic and epigenomic features that best correlate to the binding of BMAL1 to DNA. The DNA shape features Electrostatic Potential, Minor Groove Width and Propeller Twist together with the histone modifications H3K27ac, H3K4me1, H3K36me3, and H3K4me3 were the features most highly predictive of DNA binding by BMAL1 across all three tissues.

## INTRODUCTION

All animals and plants have a robust time-keeping mechanism which enables them to anticipate and adapt to periodic changes in the environment. In mammals, this time keeping mechanism, also known as the circadian system, is made up of a hierarchy of oscillators. A central clock in the suprachiasmatic nucleus (SCN) of the hypothalamus coordinates peripheral clocks in multiple tissues (Ko & Takahashi, 2006). The intracellular gene regulatory network of both the central and peripheral circadian clocks involves a relatively small set of master transcription factors (TFs) connected through negative and positive feedback loops (Takahashi et al., 2008). The core activators of the circadian network, the Clock Locomotor Output Cycles Kaput (CLOCK) and Brain and Muscle ARNT Like 1 (BMAL1), form a heterodimer complex CLOCK-BMAL1. In the classical model of clock gene regulation, the CLOCK-BMAL1 complex binds to a consensus hexanucleotide sequence known as the E-box motif (CANNTG) within the promoter or enhancer regions of clock-controlled genes to regulate their transcription (Cox & Takahashi, 2019). Alterations in the expression or binding activity of the core clock TFs disrupt natural circadian oscillations, and can lead to numerous pathologies, including insomnia, cancer, cardiovascular disease, and metabolic disorder (Kathiresan & Srivastava, 2012; Schödel et al., 2012). Here we attempt to improve our understanding of gene regulation by the CLOCK-BMAL1 complex and its perturbation by developing interpretable predictive models of DNA binding by BMAL1. Genome-wide identification of transcription factor binding sites (TFBS) is a challenging problem. Typically, only a small fraction of classically defined sequence motifs for a particular TF are bound (Lambert et al., 2018). For example, the canonical E-box binding motif for the CLOCK-BMAL1 heterodimer occurs more than 7 million times across the mouse genome, with less than 0.7% of these motifs bound by CLOCK-BMAL1 in mouse liver (Beytebiere et al., 2019). The binding of a particular TF to its cognate DNA motif depends on several molecular features including the core DNA sequence motif, flanking sequences, chromatin accessibility, local shape of the DNA, presence of collaborative or competitive co-factors, histone modifications, DNA methylation, and biophysical parameters (Dror et al., 2015; Morgunova & Taipale, 2017; J. Wang et al., 2012; T. Zhou et al., 2015). These features and their relative contribution to binding vary across cell and tissue type. Chromatin immunoprecipitation followed by massively parallel sequencing (ChIP-Seq) is the current gold standard for assaying genome-wide TF binding locations. However, each ChIP-Seq experiment measures a specific protein in a given cell type under a particular treatment condition and is dependent on availability of ChIP-grade TF-specific antibodies. Assaying the binding of a given TF under various conditions and in different tissues can thus be prohibitively expensive. As such, in recent years a number of predictive computational models of genome-wide TF binding have been developed. From these studies, DNA sequence and chromatin accessibility emerge as the most important determinants of TF binding patterns (Arvey et al., 2012; Gordân et al., 2013; Pique-Regi et al., 2011). Chromatin accessibility assays such as deoxyribonuclease hyper-sensitive sites sequencing (DNase-seq), formaldehyde-assisted isolation of regulatory elements sequencing (FAIRE-seq) and assay for transposase-accessible chromatin sequencing (ATAC-seq) have been used to improve TFBS prediction (Das & Dai, 2007). Recently, improved model predictions for TF binding have been obtained by leveraging advancements in machine learning and particularly deep learning techniques (Alipanahi et al., 2015; Quang & Xie, 2016; J. Zhou & Troyanskaya, 2015).

In this study, we present an interpretable machine learning-based models capable of predicting which canonical E-box motifs occurring in accessible chromatin regions in mouse liver, heart, and kidney bind BMAL1. Using this model, we identified genomic and epigenomic features most predictive of DNA binding by BMAL1. Our predictive model makes use of the XGBoost machine learning algorithm (Chen & Guestrin, 2016), with logistic regression used as a baseline algorithm to evaluate model performance. Published data from a ChIP-Seq study (Beytebiere et al., 2019) was used to train and test the model. Analysis of model results showed that the combination of the core E-box sequence along with two proximal flanking nucleotides, DNA shape features, and histone modifications are key determinants of BMAL1 binding across the three mouse tissues. These findings add to a previous study showing that binding specificity of the yeast basic-helix-loop-helix (bHLH) TFs Cbf1 and Tye7 to E-box motifs is governed by sequences flanking the E-box through modification of DNA shape (Grove et al., 2009). DNA shape (Mathelier et al., 2016) and histone modifications (Y. Wang et al., 2014)have been shown to be efficient predictors of TF binding in addition to DNA sequence. We investigated the role of the two central nucleotides in the core E-Box motif CANNTG in DNA binding by BMAL1. We found that the nucleotide-pair **CG** was over-represented at the center of the BMAL1-bound E-box binding motifs. The occurrence of a G in the third flanking nucleotide position upstream of the core E-box motif increased the likelihood of DNA binding by BMAL1 and was over-represented in BMAL1-bound regions in the liver. Otherwise, most of the flanking DNA sequence features showed little to no importance in predicting the binding of BMAL1 to the E-box. On the other hand, the histone modifications H3K27ac, H3K4me1, H3K36me3 and DNA shape features Electrostatic Potential (EP), ROLL, and Minor Groove Width (MGW) were efficient predictors of DNA binding by BMAL1.

## METHODS

### ChIP-Seq dataset preprocessing

Uniformly processed BMAL1 ChIP-Seq peaks from the C57BL/6J mouse liver, kidney and heart were obtained from Gene Expression Omnibus under the accession code GSE110604 (Beytebiere et al., 2019). The locations of accessible regions in DNase I-hypersensitive (DHS) sites for all three tissues (DNase-Seq) were downloaded from the Encyclopedia of DNA Elements (ENCODE, http://genome.ucsc.edu/ENCODE/). The Genome Reference Consortium Mouse Build 38 (GRCm 38) of the C57BL/6J mouse was used as the reference genome. DHS sequences were processed in Python to extract all E-Box sequences (CANNTG). To obtain all the BMAL1 ChIP-Seq peaks in accessible chromatin regions, we intersected each tissue DNase-Seq bed file with the respective BMAL1 ChIP-Seq bed file using BEDTools (Quinlan & Hall, 2010). Overlapping motifs from DNase-Seq and BMAL1 ChIP-Seq peaks in each tissue were extracted. E-box motifs in accessible chromatin regions but not overlapping with the respective tissue ChIP-Seq bed files were preprocessed and used as instances of unbound motifs (the negative dataset for the model). There were multiple instances in all three mouse tissues where there was more than one E-box motif under a BMAL1 ChIP-Seq peak. We then extracted all singleton E-boxes (i.e. instances of only one E-box motif under a BMAL1 peak) and E-boxes that were closest to the summit of the BMAL1 ChIP-Seq peaks for non-singleton E-boxes. These E-boxes were labeled as bound (the positive dataset). We extended each of the E-box motif sequences to include 4bp flanking sequences upstream and downstream. Since the E-box motif sequence is a palindrome, the reverse complement was ignored. Each E-box, thus represented by a 14-nucleotide sequence (6 bp core plus 4 bp on either end), was one-hot encoded: 1000 for adenine, 0100 for cytosine, 0010 for guanine, and 0001 for thymine. The binary (bound and unbound) E-box data collected as above produced a highly imbalanced dataset, as there are far more unbound than bound E-boxes in the mouse genome. Specifically, negative samples outnumbered positive samples by factors of 51 in the liver, 223 in the heart, and 82 in the kidney.

### DNA Shape Preprocessing

Because of the degrees of freedom of the DNA sugar phosphate backbone, neighboring base pairs and bases within a pair can vary their position relative to each other causing a change in the shape of the DNA either through rotation or translation. It has been proposed that in addition to recognizing particular DNA sequences, TFs have a preference for specific DNA conformations (Slattery et al., 2014). Incorporation of DNA shape features led to improved prediction of in vivo binding of TFs from the basic helix–loop–helix (bHLH) family (Mathelier et al., 2016). Specifically, five distinct shape features: Electrostatic Potential (EP), Minor Groove Width (MGW), Propeller Twist (ProT), Roll, and Helix Twist (HelT) have been shown to be significant for TF-DNA binding prediction (J. Li et al., 2017). We used the R/Bioconductor package DNAshapeR (Chiu et al., 2016) to evaluate DNA shape features. The DNAshapeR algorithm predicts DNA shape from DNA sequence and encodes shape feature vectors which we added to sequence-based models to predict DNA binding of BMAL1. The feature vectors for each shape category were then normalized to values between 0 and 1 by the Min-Max normalization method and placed in bins of 10bp for MGW, PROT and EP and bins of 11bp for HelT and Roll to be used as inputs for the predictive model. The number of bins for each shape feature is based on the feature matrix generated by encoding the DNA sequence.

### Histone Modification Preprocessing

We downloaded ChIP-seq data for five histone (H3) modifications, namely H3K27, H3K41, H3K43, H3K273 and H3K363, for mouse liver, kidney and heart tissues from ENCODE. These histone modification features were chosen were chosen based on data availability for all tissues and their roles in transcription factor binding explained in literature. The core E-Box sequence motifs were extended to form a 2kb region centered at the motif. The corresponding bed file was used to compute the scores for gene, promoter and enhancer regions and to generate profiles and heatmaps using deepTools (Ramírez et al., 2016). From the profiles and heatmaps generated, we found the histone modification Chip signals to extend to at most 1.5kb centered at the E-box core motif. Using the 1.5kb region centered at the E-box core motif, we extracted the histone modification features for the binary dataset for each tissue using bwtool (Pohl & Beato, 2014). The features were then divided into 10 bins with the same number of samples in each bin. This resulted in 24 features used as input for each E-box motif: 14 sequence features (one for each nucleotide), 5 DNA-Shape features and 5 histone modification features.

### Machine Learning Models

#### XGBoost

Extreme Gradient Boosting (XGBoost) is an ensemble learning method based on boosting trees for classification and regression (Chen & Guestrin, 2016). The boosted trees are added by optimizing the loss function from the previous trees, perform feature splitting to grow another tree and learn a new function to fit the residuals of the last prediction. The conventional tree boosting method uses only the first derivative of the dataset while XGBoost performs a second order Taylor expansion on the loss function leading to improved model performance (W. Li et al., 2019). From the Scikit-learn library, we performed hyperparameters tuning to adjust the following parameters to reduce the degree of overfitting and improve the accuracy of the model: the number of iterations in training the dataset (*n_estimators*), the sum of sample weight of the smallest leaf nodes to prevent overfitting (*min_child_weight*), the maximum depth of the tree in building a model for the training dataset (*max_depth*), the sampling rate of the training set in each iteration in the training samples (*subsample*), the learning rate of the training dataset (*learning_rate*), the feature sampling rate when constructing each tree (*colsample_bytree*) and the control feature to balance the positive and the negative classes in the dataset (*scale_pos_weight*).

#### Logistic Regression

Logistic regression is a parametric classification model that calculates the probability that the output variable belongs to the appropriate class (Peng et al., 2010). Logistic regression is used as the baseline for most machine learning-based classification models. In this study, we used the following logistic regression model parameters to improve performance and to reduce overfitting in our testing dataset: the regularization solver for the training dataset (*solver*), the maximum number of times the solver algorithm is run before it converges (*max_iter*), and the parameter to balance the positive and the negative classes in our dataset to have the same amount of weight (*class_weight*). These inbuilt parameters were tuned to reduce the degree of overfitting and also improve the accuracy of the model.

## RESULTS

### BMAL1 binds most frequently across tissues to the canonical E-box motif

We used the BMAL1 ChIP-Seq dataset (Beytebiere et al., 2019) and downloaded uniformly processed DNase I-hypersensitive site (DNase-seq) datasets (ENCODE) for mouse liver, heart, and kidney. For each DNase-seq dataset, the mouse mm10 reference genome was scanned and the intersection with the uniformly processed DNase-seq tracks was used to identify all instances of the E-Box motif **CANNTG**. The reverse complement of the E-box motif sequence was ignored in our scan because of its palindromic nature. The bed files processed from each tissue using the DNase-seq data were then intersected with their tissue matched ChIP-Seq dataset to extract all BMAL1 E-Box motifs (**CANNTG**) bound in open chromatin. Flanking nucleotides have been shown to affect the binding specificity of a particular TF to its cognate sequence motif (Gordân et al., 2013). We extended the E-box binding motif sequence to include 4bp upstream and downstream to identify the contribution of these flanking sequences to DNA binding by BMAL1.

Comparison of DNase-Seq signals (Abascal et al., 2020) with that of the BMAL1 ChIP-Seq signal (Beytebiere et al., 2019) revealed the DNase-seq signals to be weaker compared to ChIP-Seq signals. This could be due to different normalization methods used in processing each dataset. We also found instances where BMAL1 ChIP-seq peaks with E-box motifs were not located in open chromatin. This was true even for some core circadian clock genes. Occurrences of BMAL1-bound E-box motifs across the liver, heart and kidney were highly tissue-specific, with only 398 motifs bound in all three tissues (Fig 1A-B). Motifs bound across all tissues were over-represented among core circadian clock genes (results not shown). We computed the distribution of E-box motifs in the mouse genome using the sequence **CANNTG** (Fig 1C). The sequences **CACATG and CATGTG** were most frequently represented in the mouse genome, comprising 17.3% of all instances. These two motifs are the reverse complement of each other, but both scored the highest representation of E-box motifs in the mouse genome. Interestingly, the canonical E-Box motif **CACGTG** occurs the fewest number of times (1.83%) among all E-boxes across the mouse genome (Fig 1C). Analysis of E-box motifs from mouse liver, kidney and heart under DNase-Seq peaks showed the palindromic motif **CAGCTG** to be the most represented across open chromatin regions in the three tissues, while the canonical E-Box motif **CACGTG** was among the three least represented motifs (Fig 1D). The three least represented E-box motifs (**CACGTG, CAATTG** and **CATATG**) in accessible chromatin across all tissues were palindromes. We then used the overlap between the BMAL1 ChIP-seq and DNase-seq peaks to compute the percentage of BMAL1-bound E-box motif in the mouse liver, kidney, and heart. The kidney and heart have more bound E-box motifs as represented by the hexamer **CANNTG** in accessible chromatin than the liver. However, the kidney and heart tissue have fewer bound E-box motifs. Also, less than 20% of all individual hexamer E-box motifs found in accessible chromatin were found to be bound in all three tissues. Here, the canonical BMAL1 E-box motif **CACGTG** was the most frequently bound across all three tissues representing 18% of the **CANNTG** motifs in accessible chromatin in the liver, 15% in the kidney and 4% in the heart (Fig 1E). We observed instances where there were zero (0), exactly one (singleton) and multi (two or more) E-box motif(s) under the BMAL1 ChIP-seq signal peak across the three tissues (Fig 1F). We then extracted all singleton E-box motifs and motifs closest to the summit of the BMAL1 peak within multi-E-box peaks and labeled these as bound (positive dataset). The E-Box binding motifs in open chromatin that did not bind BMAL1 were labelled as unbound (negative dataset). The ratio of the positive to negative datasets were 1:51 in the liver, 1:82 in the kidney, and 1:223 in the heart.

**Figure 1.**
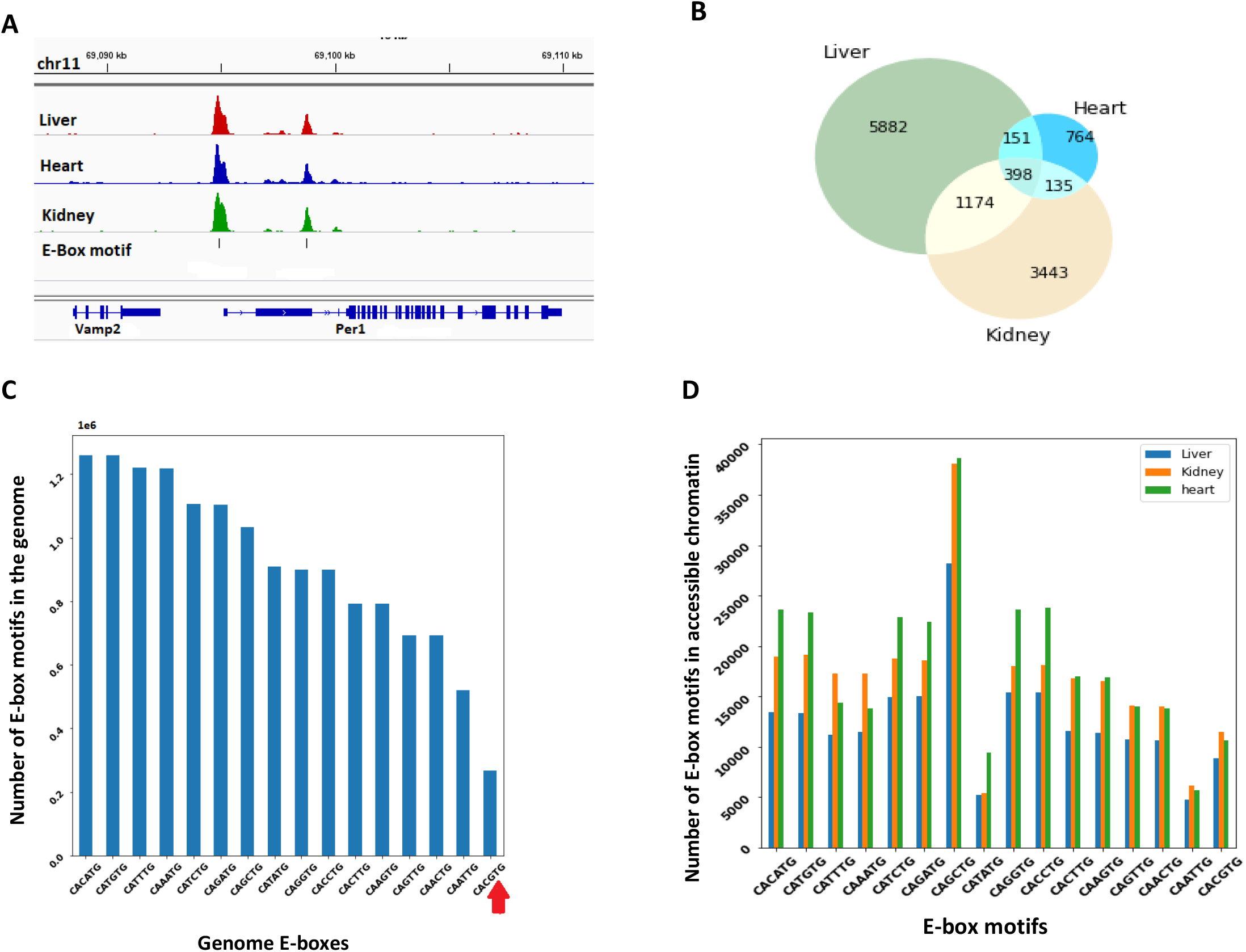
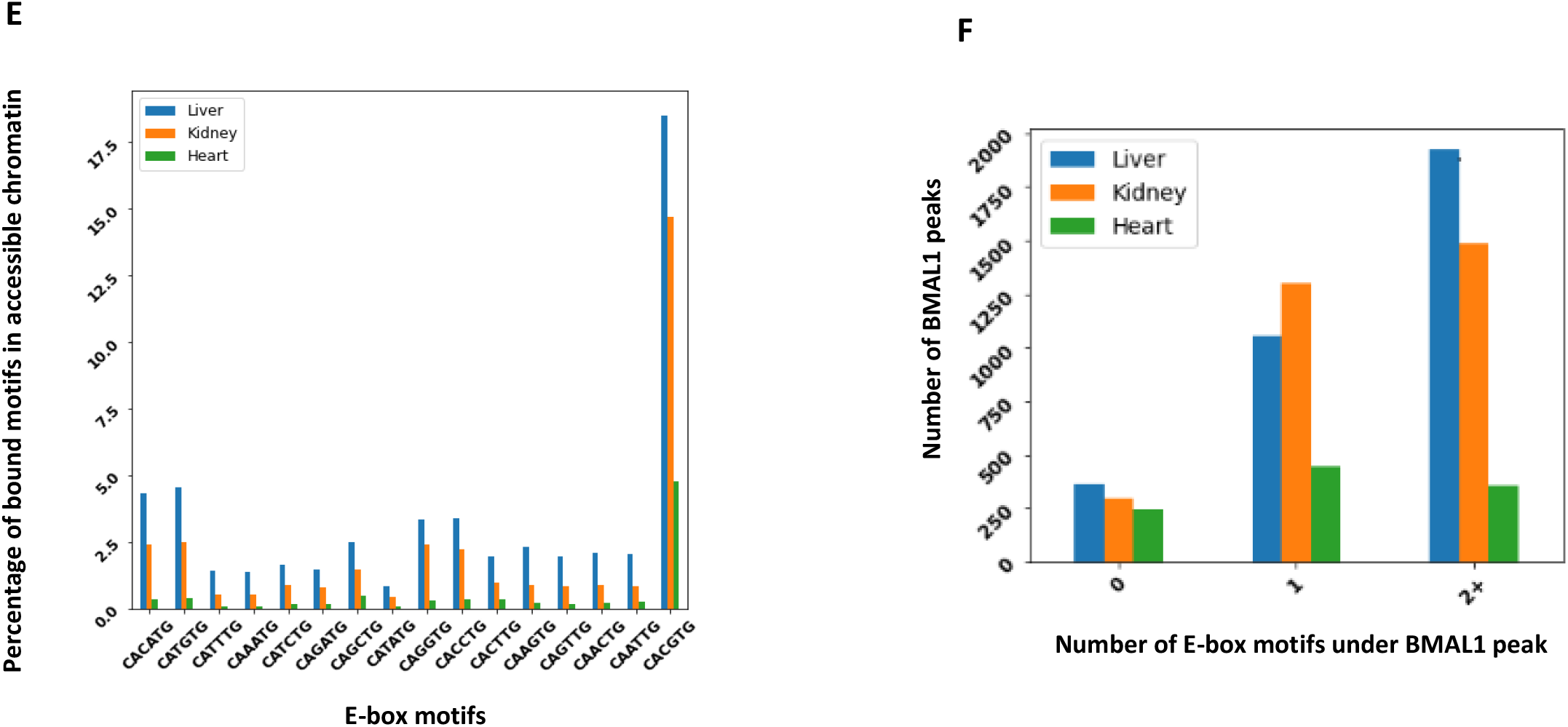
Binding of BMAL1 to E-Box motifs across multiple mouse tissues. **(A)** BMAL1 ChIP-Seq peaks in the liver (red), kidney (blue), and heart (green), and E-box binding motifs (black vertical bars) under the peaks at the Per1 locus. **(B)** Venn diagram representing the overlap of bound E-boxes in open chromatin across liver, kidney and heart. **(C)** E-box binding motif distribution across the entire mouse genome. The canonical E-Box motif CACGTG is the least represented motif in the mouse genome. **(D)** Distribution of E-box binding motifs in open chromatin across the liver (blue), kidney (orange) and heart (green). **(E)** Percentage of BMAL1 bound E-box motifs in open chromatin across the liver (blue), kidney (orange) and heart (green). **(F)** Distribution of BMAL1 peaks with zero (0-E-Box), exactly one (singleton E-box) and multi (two or more E-box) E-box motifs in the liver (blue), kidney (orange) and heart (green).

Together, these results indicate that the BMAL1 transcription factor interacts with multiple E-box motifs across the liver, kidney, and heart, with the canonical motif **CACGTG** being the most highly associated with BMAL1 binding. Also, E-box motifs are bound in a tissue-specific manner and only some core circadian clock genes are BMAL1-bound (results not shown) across all three mouse tissues studied.

### Predicting genome wide BMAL1 binding across tissues

The genomic sequence for BMAL1-bound (positive) and unbound (negative) E-box motifs was one hot-encoded (Fig 2A). Adding four flanking nucleotides upstream and downstream to the core E-box binding motif, we computed the following DNA shape features: Electrostatic Potential (EP), Minor Groove Width (MGW), Propeller Twist (ProT), Roll, and Helix Twist (HelT), using the k-mer + k-shape (k=1) sequence feature model (T. Zhou et al., 2015) (Fig 2A). Visualization of the DNA shape features EP, ProT and Roll showed significant differences in DNA shape between the bound and unbound motifs across liver, kidney, and heart, while the MGW feature showed a significant difference between the bound and the unbound motifs for the kidney only (Supplementary Figures 5&6). The shape feature vector for each shape category was then normalized to values between 0 and 1 by the Min-Max normalization method and binned in groups of 10 nucleotide for the DNA shape features EP, MGW and ProT, and groups of 12 nucleotide for HelT and Roll based on their sliding pentamer window used in computing the respective DNA shape feature. The normalized DNA shape feature vectors were then added to the encoded sequence as input features for the predictive model. Epigenetic modifications are also known to influence transcription factor binding. Specifically, histone modifications are involved in regulation of transcription factor occupancy and subsequent gene expression (Liu et al., 2017; Mathelier et al., 2016). Sequence features from the genomic regions of ± 1.5kb centered on the core E-box motif were used to compute feature vectors for five histone modifications: H3K27ac, H3K4me1, H3K4me3, H3K27me3 and H3K36me3. The ± 1.5kb region was chosen to consider histone modifications binding in the proximal promoter. The histone feature vector was then binned into 10 groups with 150bp in each bin centered on the E-box motif (Fig 2A). We implemented three different models using the final encoded feature set: (i) a DNA sequence-only model; (ii) a model based on DNA sequence and DNA shape (sequence + shape); and finally (iii) a DNA sequence, DNA shape, and histone modification (sequence + shape + HM) model. We employed two machine learning algorithms to predict which canonical E-box motifs occurring in accessible chromatin regions are bound by BMAL1 in the mouse liver, heart, and kidney. XGBoost (Chen & Guestrin, 2016) was used as our principal predictive algorithm and its performance compared with that of a baseline logistic regression model. Using grid search and stratified 5-fold cross validation, we tuned model parameters to derive the hyperparameters for each model using the liver, heart, and kidney dataset. Performance metrics from the XGBoost model outperformed that of logistic regression (Supplementary figures 1&2). The hyperparameters from the XGBoost models were used to predict the binding of BMAL1 to the E-box motif in the liver, heart and kidney (Fig 2B). The performance of the models was evaluated using the performance metrics, area under the receiver operating characteristic (AUROC) and area under the precision-recall curve AUPRC (Fig 2C)

**Figure 2:**
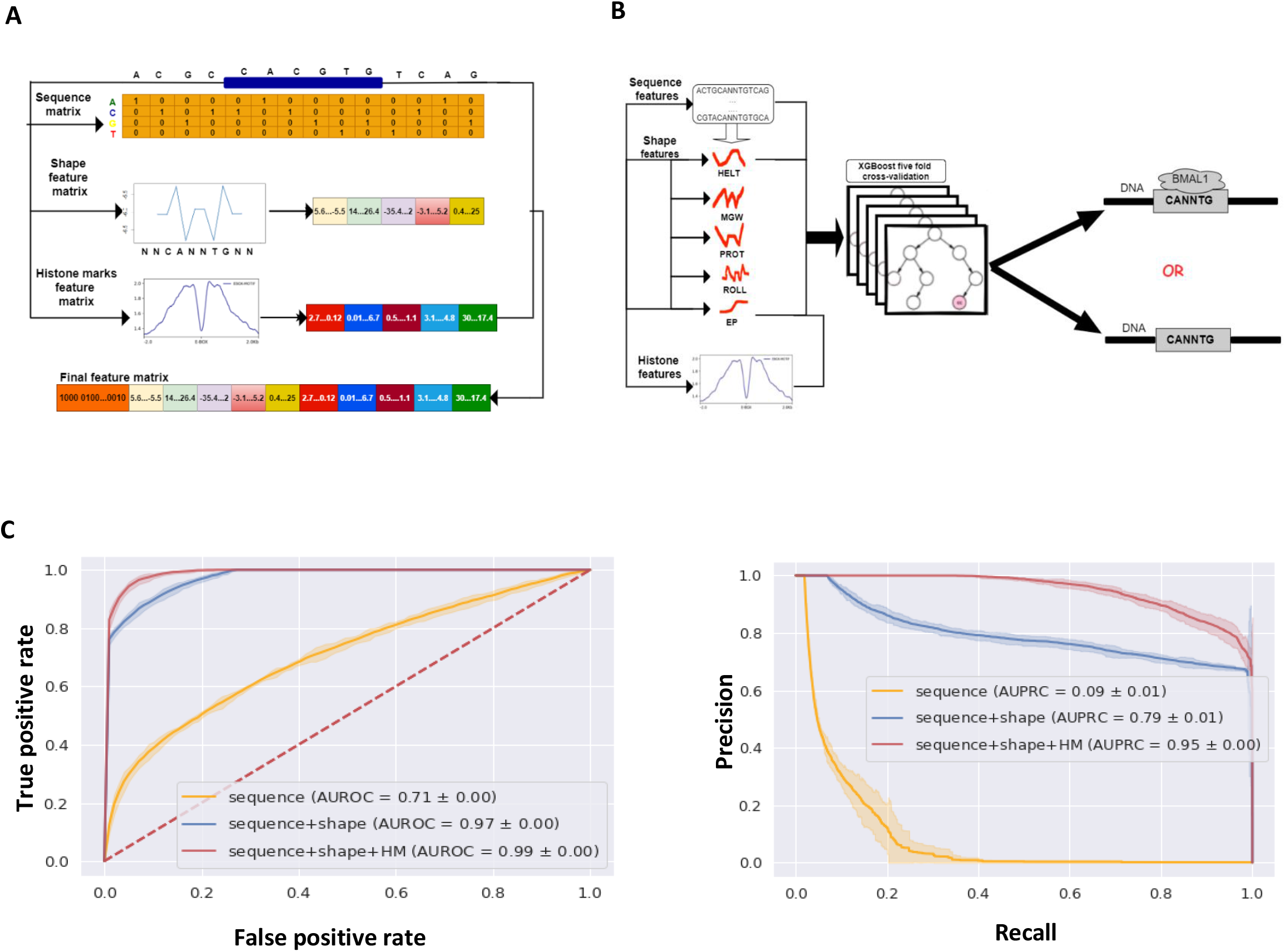
Machine learning model predicting BMAL1 binding to E-box motifs across three mouse tissues. **(A)** Design of the machine learning algorithm input features. The local chromatin features (E-box DNA sequence features) and flanking sequences were one-hot encoded. The DNA shape genomic feature matrix from the k-mer + k-shape (k=1) sequence feature model and epigenomic (histone modification) features averaged and binned were used as the final feature matrix for the model. **(B)** Schematic of the machine learning-predictive model. Based on 5-fold cross-validation, the XGBoost classifier predicted the binding status of E-box motifs in open chromatin, training on all accessible bound E-boxes and unbound E-boxes. **(C)** Performance of models predicting the binding status of E-boxes in open chromatin of the liver. The performance of each model is represented as a mean line with a shaded 95% confidence interval from 5-fold cross-validation. The legend shows the list of features used an area under the curve. Both receiver operating characteristic (ROC) and precision-recall (PRC) curves showed progressive improvement in model prediction with addition of genomic and epigenomic features.

### Within tissue (liver, kidney and heart) model performance

#### Performance of the sequence-only model

Transcription factors bind to specific DNA motifs, short DNA sequences of length 5-12bp found in promoter and enhancer regions, to regulate the expression of target genes(Ptashne, 2014; Ptashne & Gann, 2002). Binding motifs along with proximal flanking sequences upstream and downstream are key regulators of transcription factor binding to the DNA. Transcription factor binding has been studied using both experimental and computational approaches (Beytebiere et al., 2019; T. Zhou et al., 2013, 2015). The latter approach includes deep learning models (e.g., DeepSEA and DANQ, (Quang & Xie, 2016; Zheng et al., 2020) relying on convolutional neural networks. We developed two machine learning models based on logistic regression (LR) and XGBoost algorithms to predict the binding of BMAL1 to E-box motifs. Using LR as a base model, we trained and validated our XGBoost classifier on the liver, heart, and kidney with 14 features comprising nucleotides of the 6-bp core E-Box motif and ±4 flanking nucleotides. We assessed the area under the receiver operator characteristics (AUROC) and the area under the precision-recall curve (AUPRC) for each tissue using stratified 5-fold cross validation. The mean AUROC score for the liver was 0.71, 0.78 for kidney and 0.80 for the heart (Fig 3A). To further investigate the performance of the model, we evaluated the area under the precision-recall curve (AUPRC), a more appropriate performance metric given the unbalanced distribution in the two classes (bound vs unbound) in the dataset across all three tissues. The mean AUPRC score for the liver was 0.09, 0.10 for the kidney and 0.06 for the heart (Fig 3B). The AUPRC score for each tissue exceeds the corresponding baseline score given by the fraction of the positive class. The baseline AUPRC score was 0.0052, 0.0082 and 0.022 for the liver, kidney, and heart respectively. The high AUROC and AUPRC scores across all tissues suggest differences in flanking sequence between BMAL1 bound and unbound E-box motifs.

**Figure 3.**
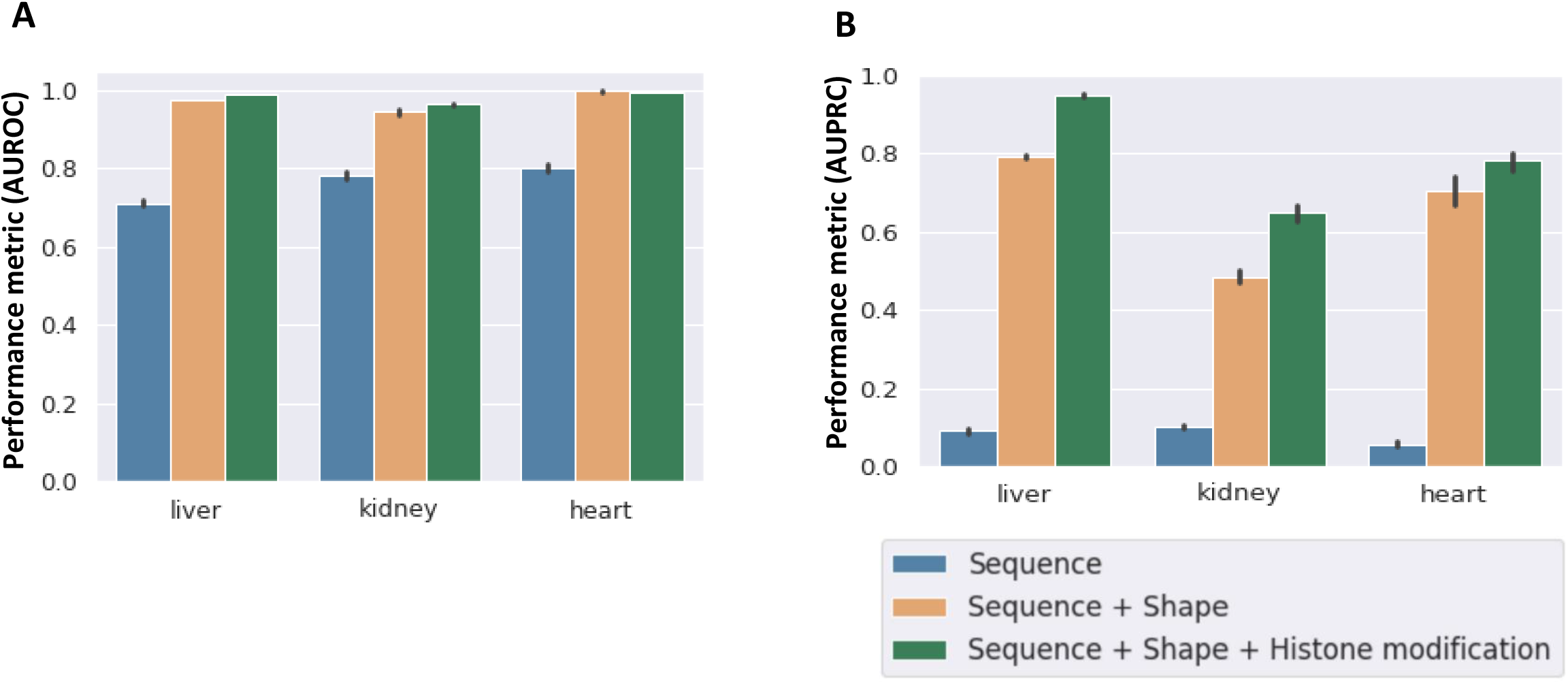
Adding DNA Shape and Histone modification (HM) features to DNA Sequence significantly improves prediction of BMAL1 binding across all tissues. **(A)** The area under the receiver operating characteristics (AUROC) for liver, kidney, and heart for the sequence-only model (blue), sequence plus DNA shape model (brown) and sequence plus DNA shape plus HM model (green). The mean AUROC increases sharply with the addition of DNA shape features to the model, with a much smaller increase associated with the addition of HMs. **(B)** The area under the precision recall curve (AUPRC) in liver, kidney and heart for the sequence-only model (blue), sequence plus DNA shape model (brown) and sequence plus DNA shape plus HM model (green). As with AUROC, the mean AUPRC increased by a large margin with the addition of DNA shape features to the model, with a smaller increase associated with the addition of HMs.

#### Adding DNA shape features to DNA sequence improves model performance

The three-dimensional structure of DNA gives rise to specific local conformations. Features to quantify DNA shape have been derived computationally using Monte Carlo (MC) simulations from local DNA sequence (J. Li et al., 2017; T. Zhou et al., 2013). Five distinct DNA shape features, Electrostatic Potential (EP), Minor Groove Width (MGW), Propeller Twist (ProT), Roll, and Helix Twist (HelT) were found to contribute to the binding affinity of transcription factors from the basic helix loop helix (bHLH) family (T. Zhou et al., 2015). Different sizes of sequence were used to compute each shape feature. The EP, MGW and ProT DNA shape features were computed from a 10bp sequence comprising the 6bp core E-box motif with ± 2bp flanking sequence, while Roll and HelT were computed from the 6bp E-box motif plus ± 3bp flanking sequence. We combined the DNA shape feature matrix with the sequence features as input for the model, to evaluate the contribution of DNA shape to BMAL1 binding. The mean AUROC performance score from five-fold cross validation model for the liver was 0.97, 0.98 for kidney and 0.98 for the heart sequence plus DNA shape dataset (Fig 3A). Compared to the sequence-only model, the mean AUPRC metric increased sharply from 0.09 to 0.79 for the liver, 0.10 to 0.51 for the kidney, and 0.06 to 0.71 for the heart (Fig 3B), suggesting significant differences in local DNA shape features between the bound and unbound E-boxes. Inspection of feature importance revealed that the EP, Roll and MGW shape features contributed 55% to prediction of BMAL1 binding to the E-box motif in the liver. For the kidney, the EP, PROT and Roll DNA shape features contributed 48% to prediction of BMAL1 binding, while in the heart, EP, ROLL and HelT contributed to 52% of the prediction. Overall, the EP, Roll and PROT DNA shape features had the biggest influence on prediction of bound E-Box motif across all three tissues (Supplementary Figure 3). A recent study (T. Zhou et al., 2015) showed that for the bHLH transcription factor Max, Roll and ProT DNA shape features were the dominant determinants of TF-DNA binding affinity. This finding agrees with our observation that the Roll and PROT features contribute to BMAL1 – E-box binding in all three tissues. Validating the predictive classifier model on only the DNA shape features did not give a convincing performance metric as compared to the DNA sequence-only model or the sequence + DNA shape model, suggesting that local shape or topology of the DNA near the E-box by itself is not sufficient to predict BMAL1 binding (results not shown). Recent studies have shown that DNA shape computed using core TF binding motifs and flanking sequences improves transcription factor binding prediction in many human TFs (Mathelier et al., 2016; S. Wang et al., 2021; T. Zhou et al., 2015). In another study, DNA topology was highly correlated with the structure and stability of the nucleosome, with topological changes influencing the binding of transcription factors to DNA (Gupta et al., 2009). These findings are consistent with our results.

#### Adding histone modifications to DNA shape and sequence gave the best model performance

Histone modifications (HMs) in gene promoter regions have been shown previously to be correlated with transcription factor binding (Benveniste et al., 2014). However, the mechanisms of interaction between transcription factor binding and HMs are not fully understood. Recent studies have shown that the extent to which HMs improve prediction of TF binding is protein-specific, with models of bHLH transcription factor binding showing significantly improved accuracy when HMs are included (Xin & Rohs, 2018a). In particular, the modifications H3K9ac, H3K79me2 and H3K4me3 are strongly associated with binding of bHLH transcription factors (Guccione et al., 2006). Based on these findings, several models have been developed to improve transcription factor binding prediction using results from epigenetic assays (Heintzman et al., 2009; Ramsey et al., 2010). We examined the importance of HMs in prediction of BMAL1 binding by adding five histone features (H3K27ac, H3K4me1, H3K4me3, H3K27me3 and H3K36me3) to the sequence and DNA shape feature matrix. These histone modification features were chosen were chosen based on data availability for all tissues and their roles in transcription factor binding explained in literature (Xin & Rohs, 2018b). The mean AUROC performance score obtained was 0.99 for the liver, 0.988 for kidney and 0.99 for the heart DNA sequence + shape + HM model (Fig 3A). The mean AUPRC performance metric increased significantly to 0.95, 0.65 and 0.79 for the liver, kidney and heart respectively (Fig 3B). The HMs with the largest contribution to BMAL1 binding were H3K27ac, H3K4me1 and H3K36me3 in both liver and heart, and H3K27ac, H3K4me1 and H3K4me3 in the kidney. These results show that the combination the transcription factor binding motif and proximal flanking sequence, local shape of the DNA, and histone modifications together impact the binding of BMAL1 to E-box motifs.

### Quantifying importance of features used in the models

Given the improved performance of the model using DNA sequence, DNA shape, and HMs, we employed the eli5 permutation importance method (Korobov & Lopuhin, 2021) to identify features most predictive of DNA binding by BMAL1 to the E-box. The feature importance for each DNA shape and histone modification feature was calculated as the sum of the feature importances of all bins for that particular feature. The feature importance of each nucleotide type at a particular position relative to the E-Box motif was normalized to the nucleotide type and the sum of all feature importances at that nucleotide position. Our analysis showed the immediate flanking sequences upstream and downstream of the core E-box binding motif were important predictors of BMAL1 binding in the liver, heart and kidney as compared to distal flanking sequence (Fig 4). Analysis of the binding specificities of the bHLH transcription factors CBf1 and Tye7 in yeast has previously shown that 2bp flanking sequences contribute to the binding of these transcription factors to the E-box (Gordân et al., 2013). In our quantitative analysis of the E-box sequence, we did not find the two central base pairs of the CANNTG E-box motif to directly contribute to the model performance across the three mouse tissues, even though BMAL1 has a strong preference for the **CG** central base pairs across all tissues given that the **CACGTG** motif was the most highly represented among BMAL1-bound E-boxes. Analysis of the feature weights showed the nucleotide **G** at the second proximal upstream flanking sequence to be a strong predictor of DNA-BMAL1 binding in the liver (Fig 4A). The nucleotide **G** at the second proximal upstream flanking sequence represented more than 50% of the feature weights used in predicting BMAL1 DNA binding in the liver. Other contributing features include the EP shape feature (10%) and the H3K27ac histone modification (6%). Most of the DNA shape and histone modification features had weights greater than 5% indicating their importance in predicting BMAL1 DNA binding in the liver, while most of the DNA sequence features except Seq-2 (the second proximal upstream flanking nucleotide), had a feature weight of less than 5%. In the kidney, the histone modification feature H3K27ac had the highest feature importance weight, contributing 21% to the overall feature importance score (Fig 4B). The shape feature EP followed with a feature weight of 19%. Three histone modifications (H3K27ac, H3K4me3 and H3K4me1) and four DNA shape features (EP, ProT and Roll) all had feature weights > 5%. In the heart, H3K27ac and H3K4me3 had the highest feature importance (all > 20%) followed by the DNA shape feature EP (8%). Most of the DNA sequence features had weights 5% in both heart and kidney. The histone modifications H3K27ac, H3K4me1, H3K4me3 and DNA shape features EP, and Roll showed high importance scores across all three tissues (Fig 4A-C). The second upstream flanking nucleotide (Seq-2) had by far the highest feature importance score in the liver. The nucleotides **G** and **C** were highly represented at the second proximal upstream flanking sequence of the liver bound E-box motifs. This was supported by the sequence logo of the bound E-box sequence with 4bp upstream and downstream of the core E-box motif (Fig 4D). The nucleotide **G** and **C**, at the second proximal flanking sequence (Seq-2) scored the highest in the liver (Fig 4A). This was not the case for the kidney and heart. Overall, the DNA shape and histone modification features H3K27ac, EP, H3K4me1, ROLL, MGW, H3K36me3, HELT, PROT and H3K4me3 had high scores across all tissues.

**Figure 4:**
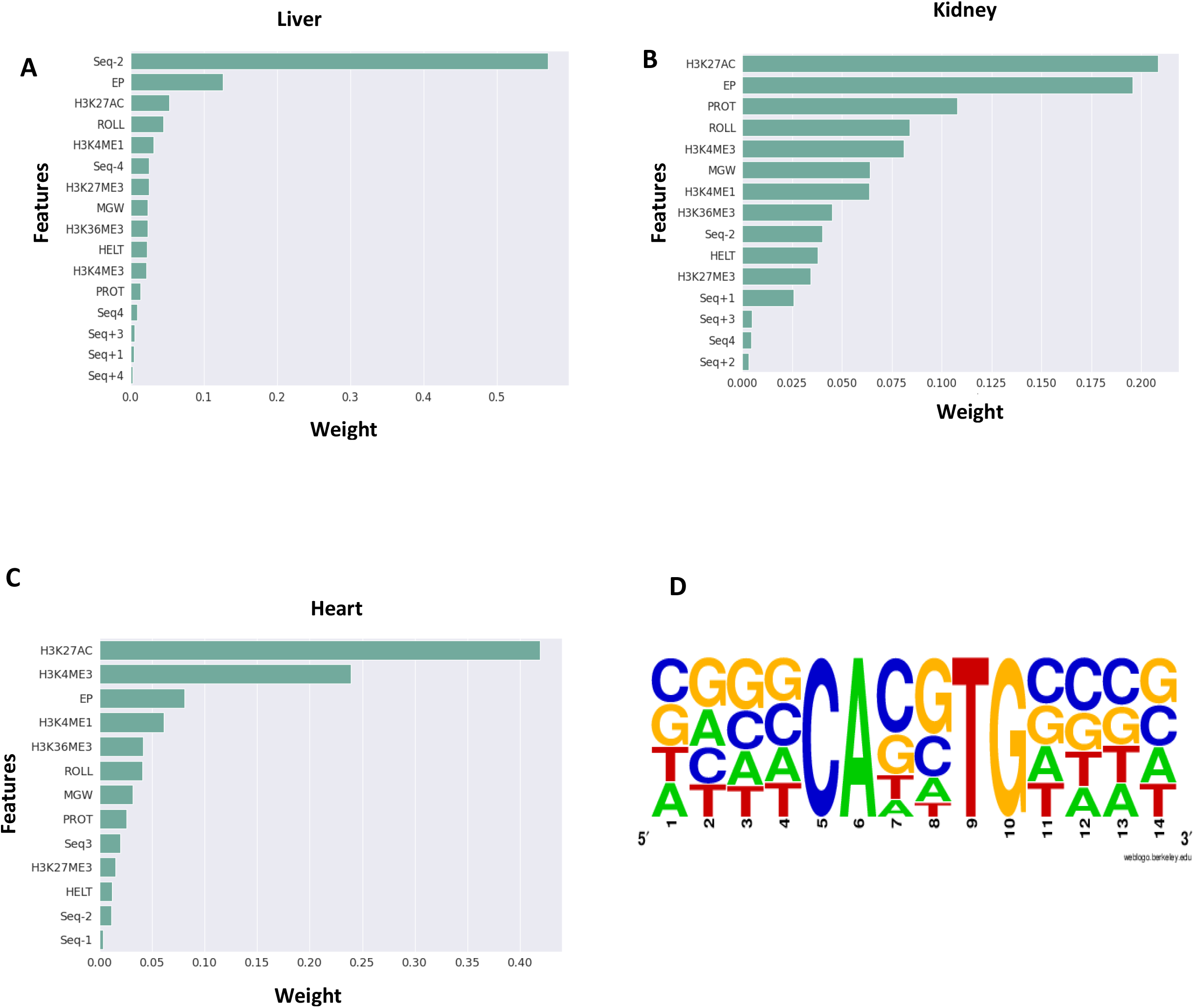
Feature importance of all genomic (sequence) and epigenomic (DNA shape and histone modification) features from the XGBoost classifier model across all tissues. Feature importance in the XGBoost classifier model in **(A)** liver, **(B)** kidney, and **(C)** heart. The feature importance for each DNA shape and histone modification feature is calculated as the sum of all the feature importance of all bins for that particular histone modification feature. The feature importance of each nucleotide type at a particular position relative to the E-Box motif is normalized to the nucleotide type and the sum of all feature importance at that nucleotide position. **(D)** Standard plot sequence logo for BMAL1 bound E-box motifs in the liver.

### DNA shape and histone modification features common to all three tissues is not a driving force for cross tissue prediction

We found 398 E-boxes in open chromatin common to all three tissues, 1174 common to the liver and kidney, 135 common to kidney and heart, and 151 common to the heart and the liver (Fig 1B). To test the hypothesis that the DNA binding of BMAL1 is determined by similar factors across the three tissues, we developed cross-tissue models for binding prediction based on (a) sequence features only; (b) sequence plus DNA shape features; and (c) sequence plus DNA shape plus histone modifications. We trained these three models using the hyper-parameters derived for within-tissue models for each tissue. The trained models were then used to predict BMAL1 binding in a different tissue. The performance of the sequence-only model was preserved across all three tissues, i.e., the AUROC, AUPRC and the true positives ratio of the cross-tissue models were similar to that of the within-tissue models (Fig 5A,blue bars). Using the sequence only model, AUROC and AUPRC scores for the models trained on the liver data and used to predict binding on the kidney dataset (liver_kidney) and trained on the kidney dataset for predicting binding in the liver (kidney_liver), were comparable to the AUROC and AUPRC scores for the liver within-tissue model (train-test on liver) and the kidney within-tissue model (train-test on kidney). Surprisingly, adding the DNA shape and HM features decreased the performance scores across all the cross-tissue models (Fig 5A-C). The AUROC and AUPRC scores for the cross-tissue models decreased with the addition of DNA shape features (Fig 5A-B, brown bars). This was reflected in the correct prediction (true positives) in one tissue when the model is trained on another tissue. The correct prediction of the bound and unbound classes in each tissue is relevant for generating a higher prediction performance score metrics (AUROC and AUPRC). The model trained on the liver was able to predict 22% of the E-boxes bound in the kidney (liver_kidney) and 22% of those bound in the heart (liver_heart) using the sequence plus DNA shape model (Fig 5C,brown bars). The model predicted most of the bound class in the kidney and heart as unbound, yielding a high false negative rate ratio. The addition of the histone modification features improved the AUROC and AUPRC model performance scores for most of the cross-tissue models (Fig 5A-B, green bars). The cross-tissue sequence plus DNA shape plus HM model trained on the liver dataset correctly predicted only 18% of E-boxes bound in the kidney and 19% of those bound in the heart, leading to a high false negative rate. The sequence-only and the sequence plus DNA shape plus HM models performed similarly when trained on the kidney and tested on the liver dataset. The AUROC performance score slightly increased from 0.68 for the sequence plus DNA shape model to 0.83 for the sequence plus DNA shape plus HM model, while the AUPRC score increased sharply from 0.055 to 0.38 for the model trained on the kidney dataset to predict binding in the liver. These results show that while the sequence-only model performed well in cross-tissue binding prediction, adding genomic features like DNA shape and epigenetic features like histone modifications highlighted the tissue specificity in binding of BMAL1 to DNA. Thus, the tissue specific binding of BMAL1 to DNA is not highly dependent on the E-box binding motifs but other genomic features like DNA shape and epigenetic features like histone modifications and likely other competitive transcription factors.

**Figure 5.**
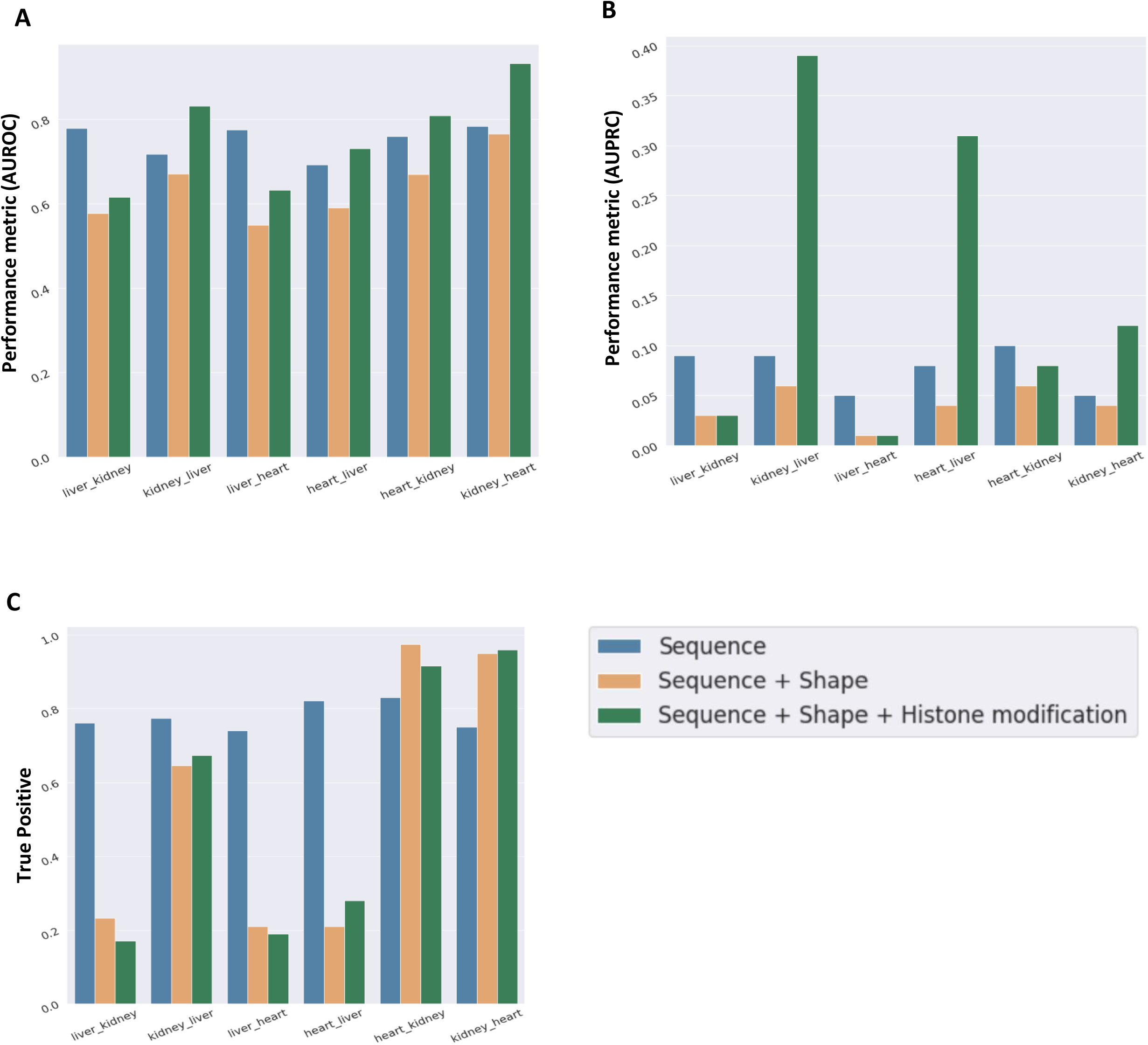
Performance metrics for cross-tissue prediction models. Scores for the liver, kidney and heart sequence-only model (blue bars), sequence plus DNA shape model (brown bars) and sequence plus DNA shape plus HM model (green bars)**: (A)** Area under the receiver operating characteristics (AUROC); **(B)** Area under the precision recall curve (AUPRC); **(C)** True positive rates. (Notation explanation: liver_kidney refers to the model trained on the liver dataset and used to predict binding on the kidney dataset.)

## Discussion

The identification of transcription factor binding determinants to DNA sequence can improve our understanding of gene regulatory grammar (Levine & Tjian, 2003). However, the precise DNA-binding sequences for most TFs are currently unknown. Here we focus on prediction of tissue-specific DNA binding by BMAL1, a master regulator of the circadian clock. While the architecture of the core clock gene regulatory network in the suprachiasmatic nucleus in the brain and other peripheral tissues is believed to be similar, clock-controlled gene expression is largely tissue-specific (Beytebiere et al., 2019; Mure et al., 2018). Here we use XGBoost, an ensemble decision tree-based machine learning algorithm, to predict the binding of BMAL1 to its putative binding motif (the E-box) in three mouse tissues (liver, heart and kidney). The AUROC and AUPRC performance metrics were used to assess the performance of the model on each tissue. The canonical E-box binding motif **CACGTG** occurs the fewest number of times in the entire genome and in accessible open chromatin, but it was the most frequently bound E-box motif across all three tissues. The heart tissue had more E-box motifs in accessible chromatin than the liver and kidney; however, it had the fewest bound E-box motifs in accessible chromatin. The role of circadian rhythms in the heart is not well understood. Only 6% of protein coding genes in the mouse heart tissue are circadian regulated as compared to 11%-16% found in the liver (Zhang et al., 2014), which could be accounted for by less genome-wide DNA binding by BMAL1. Analysis of our predictive model did not show the two middle base pairs in the generic E-box motif **CANNTG** to directly influence the DNA-binding preference of BMAL1. We employed three different models: sequence-only, sequence plus DNA shape, and sequence plus DNA shape plus histone modification, for binding prediction. Our interpretable models showed that the ability to predict binding of BMAL1 to DNA improved when genomic (DNA shape) and epigenomic features (histone modifications) were added to sequence-only models. The sequence plus DNA shape plus histone modification model generated the highest performance score across all three tissues. Analysis of feature importance showed the histone modification H3K27ac, and the DNA shape feature Electrostatic Potential dominated binding prediction across the kidney and the heart tissue. DNA sequence features by themselves had little to no effect on binding prediction in the kidney and heart. However, the second nucleotide upstream of the E-box had a large contribution to BMAL1-DNA binding in the liver. The nucleotide **G** in this position had the highest weight, contributing to about 50% of the feature importance score. Based on these results, we propose that the local shape of the DNA and histone modification features are the key driving factors in BMAL1 binding to the E-box.

Several experimental models have shown the tissue specificity or the more accessibility of transcription factor binding in specific tissues (Beytebiere et al., 2019; Mure et al., 2018). We investigated tissue specific BMAL1 binding by building cross tissue models trained on one tissue and tested in another tissue with the same set of features. The sequence-only model performance was preserved for between-tissue prediction with AUROC and AUPRC performance scores comparable to the within-tissue models. Adding DNA shape and histone modification features to DNA sequence gave a poor model performance score for the cross-tissue models. These results are consistent with tissue specific DNA binding of BMAL1 to differentially regulate gene expression. Our model shows that the tissue specificity of BMAL1-DNA binding is not explained when considering only DNA sequence. Additional features (DNA shape and histone modification) are necessary to account for the tissue specificity of DNA binding by BMAL1. Thus, the model performance scores from training the model on one tissue and predicting BMAL1-DNA binding in other tissue were lower than the within tissue models. In summary, our results provide novel insights into binding of BMAL1 to E-box motifs to regulate rhythmic gene expression in a tissue-specific manner. Our models also revealed the importance of genomic and epigenomic features in BMAL1-DNA binding. These findings are likely to improve our understanding of how transcription factors bind to their cognate motifs to regulate tissue-specific gene expression.

## Supplementary figure captions

Supplementary figure 1: **Model performance scores. (A, C)** Performance of models predicting the binding status of E-box in open chromatin of the heart using XGBoost. Performance of each model is represented as a mean line with a 95% confidence interval shaded around the line resulting from 5-fold cross validation. The legend shows the list of features used, as well as area under the curve. Both receiver operating characteristic (ROC) and precision recall (PRC) curves are shown. **(B, D)** Performance of models predicting the binding status of E-box in open chromatin of the heart using logistic regression. Performance of each model is represented as a mean line with a 95% confidence interval shaded around the line resulting from 5-fold cross validation. The legend shows the list of features used, as well as area under the curve. Both receiver operating characteristic (ROC) and precision recall (PRC) curves are shown.

Supplementary figure 2: **Model performance scores. (A, C)** Performance of models predicting the binding status of E-box in open chromatin of the kidney using XGBoost. Performance of each model is represented as a mean line with a 95% confidence interval shaded around the line resulting from 5-fold cross validation. The legend shows the list of features used, as well as area under the curve. Both receiver operating characteristic (ROC) and precision recall (PRC) curves are shown. **(B, D)** Performance of models predicting the binding status of E-box in open chromatin of the kidney using logistic regression. Performance of each model is represented as a mean line with a 95% confidence interval shaded around the line resulting from 5-fold cross validation. The legend shows the list of features used, as well as area under the curve. Both receiver operating characteristic (ROC) and precision recall (PRC) curves are shown.

Supplementary figure 3: **Feature importance of all genomic (sequence and DNA shape) features from the XGBoost classifier model across all tissues.** Feature importance in the XGBoost classifier model in **(A)** liver, **(B)** kidney, and **(C)** heart. The feature importance for each DNA shape feature is calculated as the sum of all the feature importance of all bins for that particular DNA shape feature. The feature importance of each nucleotide type at a particular position relative to the E-Box motif is normalized to the nucleotide type and the sum of all feature importance at that nucleotide position.

Supplementary Figure 4: **Liver DNA Shape features.** Electrostatic Potential (EP) at **(A)** BMAL1 bound (**B)** BMAL1 unbound E-boxes. Roll at **(C)** BMAL1 bound **(D)** BMAL1 unbound E-boxes. Propeller Twist (ProT) at **(E)** BMAL1 bound **(F)** BMAL1 unbound E-boxes. The differences in the bound and unbound shape features contribute to the model accurately predicting BMAL1 binding to the E-box motifs. The mean values represent the number of E-box motifs used in generating the shape figure.

Supplementary Figure 5: **Heart DNA Shape features.** Electrostatic Potential (EP) at **(A)** BMAL1 bound (**B)** BMAL1 unbound E-boxes. Roll at **(C)** BMAL1 bound **(D)** BMAL1 unbound E-boxes. Propeller Twist (ProT) at **(E)** BMAL1 bound **(F)** BMAL1 unbound E-boxes. The differences in the bound and unbound shape features contribute to the model accurately predicting BMAL1 binding to the E-box motifs. The mean values represent the number of E-box motifs used in generating the shape figure.

Supplementary Figure 6: **Kidney DNA Shape features.** Electrostatic Potential (EP) at **(A)** BMAL1 bound (**B)** BMAL1 unbound E-boxes. Roll at **(C)** BMAL1 bound **(D)** BMAL1 unbound E-boxes. Propeller Twist (ProT) at **(E)** BMAL1 bound **(F)** BMAL1 unbound E-boxes. The differences in the bound and unbound shape features contribute to the model accurately predicting BMAL1 binding to the E-box motifs. The mean values represent the number of E-box motifs used in generating the shape figure.

Supplementary Figure 7: **Heart histone modification profiles.** H3K27ac at **(A)** BMAL1 bound **(B)** BMAL1 unbound E-boxes. H3K4ME1 at **(C)** BMAL1 bound **(D)** BMAL1 unbound E-boxes. H3K4ME3 at **(E)** BMAL1 bound **(F)** BMAL1 unbound E-boxes. The differences in the bound and unbound profile shapes contribute to the model accurately predicting BMAL1 binding to the E-box motifs.

Supplementary Figure 8: **Heart histone modification profiles.** H3K27ac at **(A)** BMAL1 bound **(B)** BMAL1 unbound E-boxes. H3K4ME1 at **(C)** BMAL1 bound **(D)** BMAL1 unbound E-boxes. H3K4ME3 at **(E)** BMAL1 bound **(F)** BMAL1 unbound E-boxes. The differences in the bound and unbound profile shapes contribute to the model accurately predicting BMAL1 binding to the E-box motifs.

Supplementary Figure 9: **Confusion matrix for cross tissue models. (A)** Liver_Heart **(B)** Liver_Kidney **(C)** Heart_Liver **(D)** Heart_Kidney **(E)** Kidney_Heart **(F)** Kidney_Liver (Liver-Heart means the model was trained on the all the liver dataset and used to predict binding in the heart dataset.)

