## Supplemental figures for "Prediction of mammalian tissue-specific CLOCK-BMAL1 binding to E-box motifs"

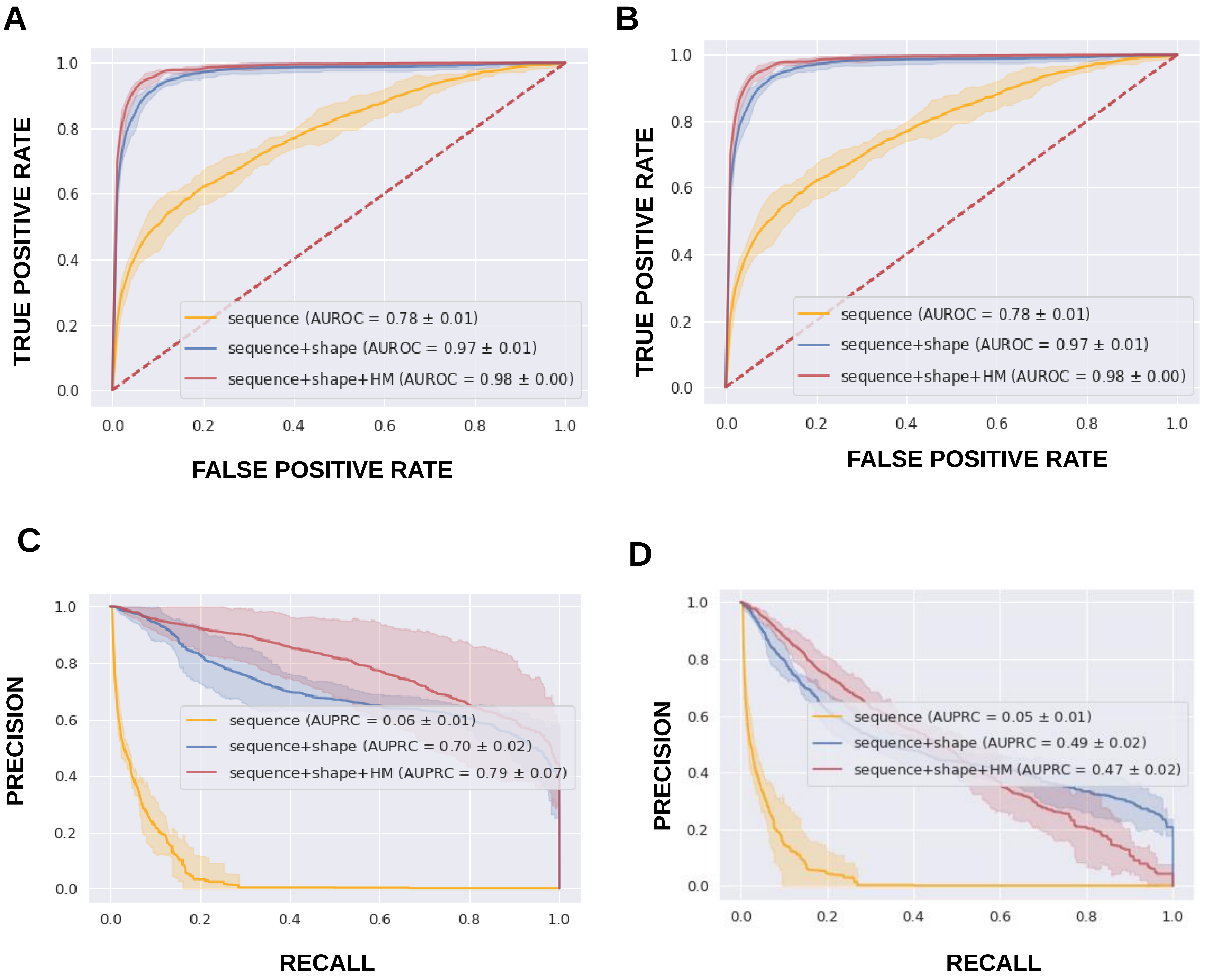


• Supplementary figure 1. **Model performance scores. (A, C)** Performance of models predicting the binding status of E-box in open chromatin of the heart using XGBoost. Performance of each model is represented as a mean line with a 95% confidence interval shaded around the line resulting from 5-fold cross validation. The legend shows the list of features used, as well as area under the curve. Both receiver operating characteristic (ROC) and precision recall (PRC) curves are shown. **(B, D)** Performance of models predicting the binding status of E-box in open chromatin of the heart using logistic regression. Performance of each model is represented as a mean line with a 95% confidence interval shaded around the line resulting from 5-fold cross validation. The legend shows the list of features used, as well as area under the curve. Both receiver operating characteristic (ROC) and precision recall (PRC) curves are shown.

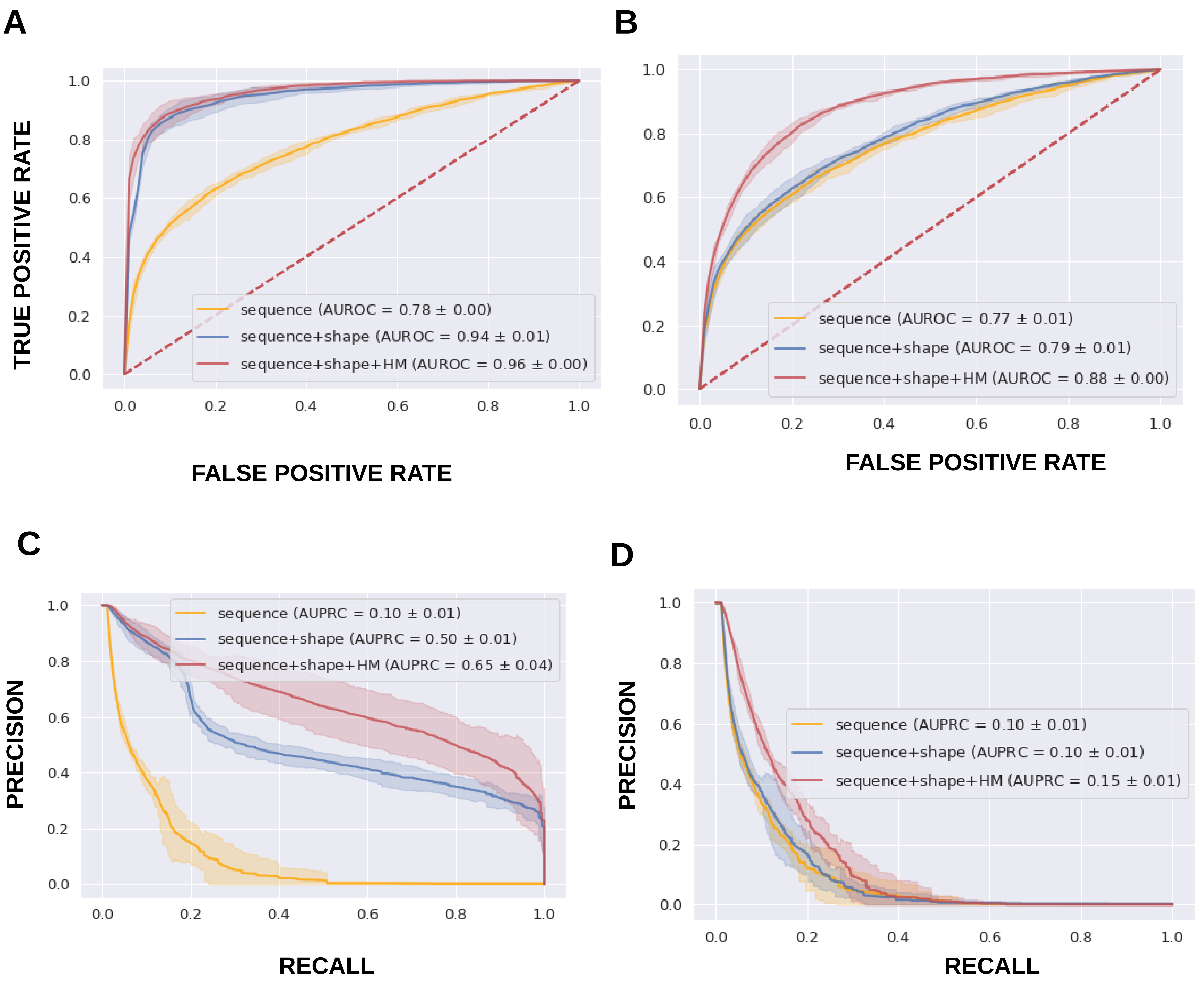


• Supplementary figure 2. **Model performance scores. (A, C)** Performance of models predicting the binding status of E-box in open chromatin of the kidney using XGBoost. Performance of each model is represented as a mean line with a 95% confidence interval shaded around the line resulting from 5-fold cross validation. The legend shows the list of features used, as well as area under the curve. Both receiver operating characteristic (ROC) and precision recall (PRC) curves are shown. **(B, D)** Performance of models predicting the binding status of E-box in open chromatin of the kidney using logistic regression. Performance of each model is represented as a mean line with a 95% confidence interval shaded around the line resulting from 5-fold cross validation. The legend shows the list of features used, as well as area under the curve. Both receiver operating characteristic (ROC) and precision recall (PRC) curves are shown.

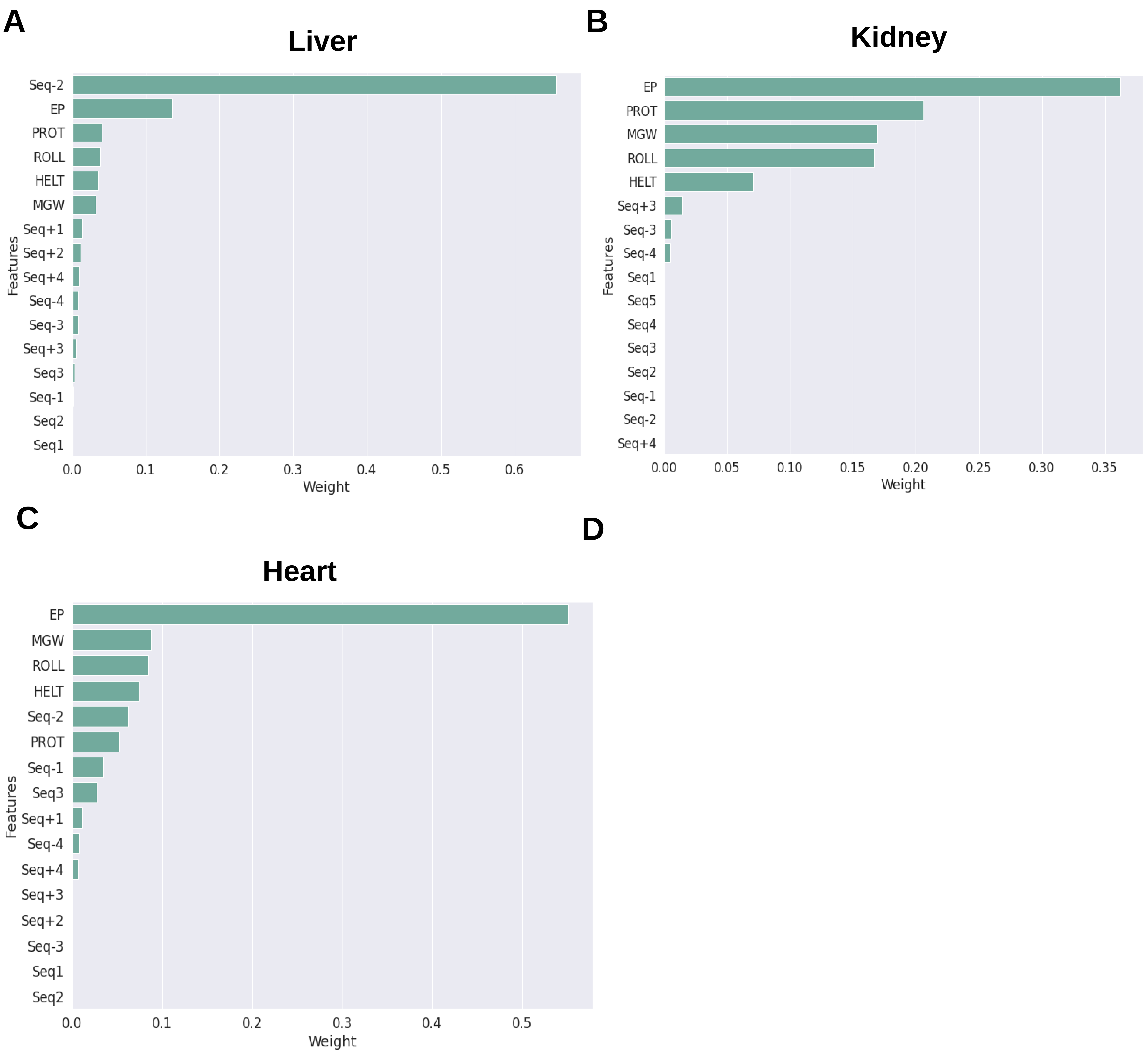


Supplementary figure 3. **Feature importance of all genomic (sequence and DNA shape) features from the XGBoost classifier model across all tissues**. Feature importance in the XGBoost classifier model in **(A)** liver, **(B)** kidney, and **(C)** heart. The feature importance for each DNA shape feature is calculated as the sum of all the feature importance of all bins for that particular DNA shape feature. The feature importance of each nucleotide type at a particular position relative to the E-Box motif is normalized to the nucleotide type and the sum of all feature importance at that nucleotide position.

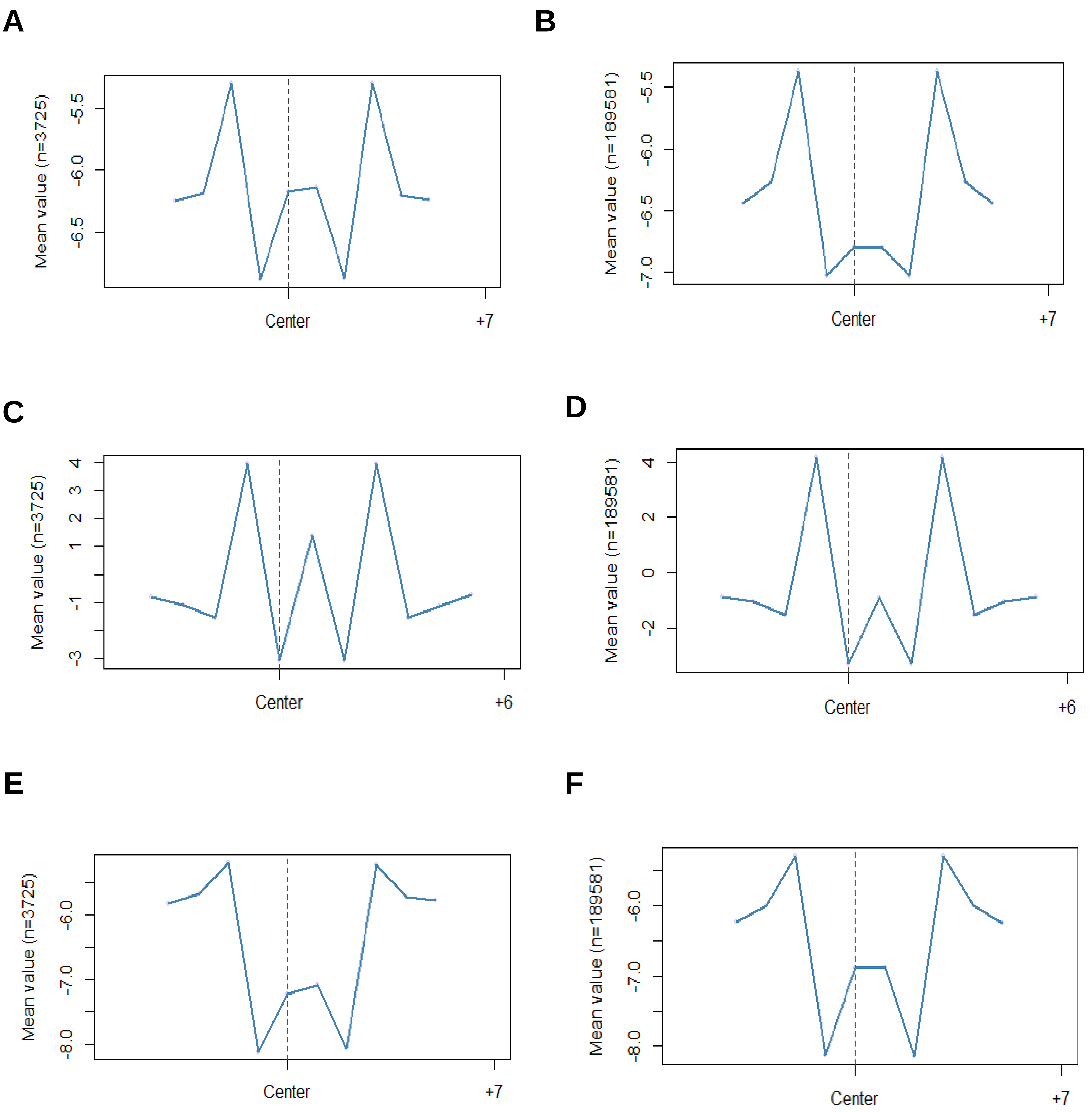


• Supplementary Figure 4: **Liver DNA Shape features.** Electrostatic Potential (EP) at **(A)** BMAL1 bound **(B)** BMAL1 unbound E-boxes. Roll at **(C)** BMAL1 bound **(D)** BMAL1 unbound E-boxes. Propeller Twist (ProT) at **(E)** BMAL1 bound **(F)** BMAL1 unbound E-boxes. The differences in the bound and unbound shape features contribute to the model accurately predicting BMAL1 binding to the E-box motifs. The mean values represent the number of E-box motifs used in generating the shape figure.

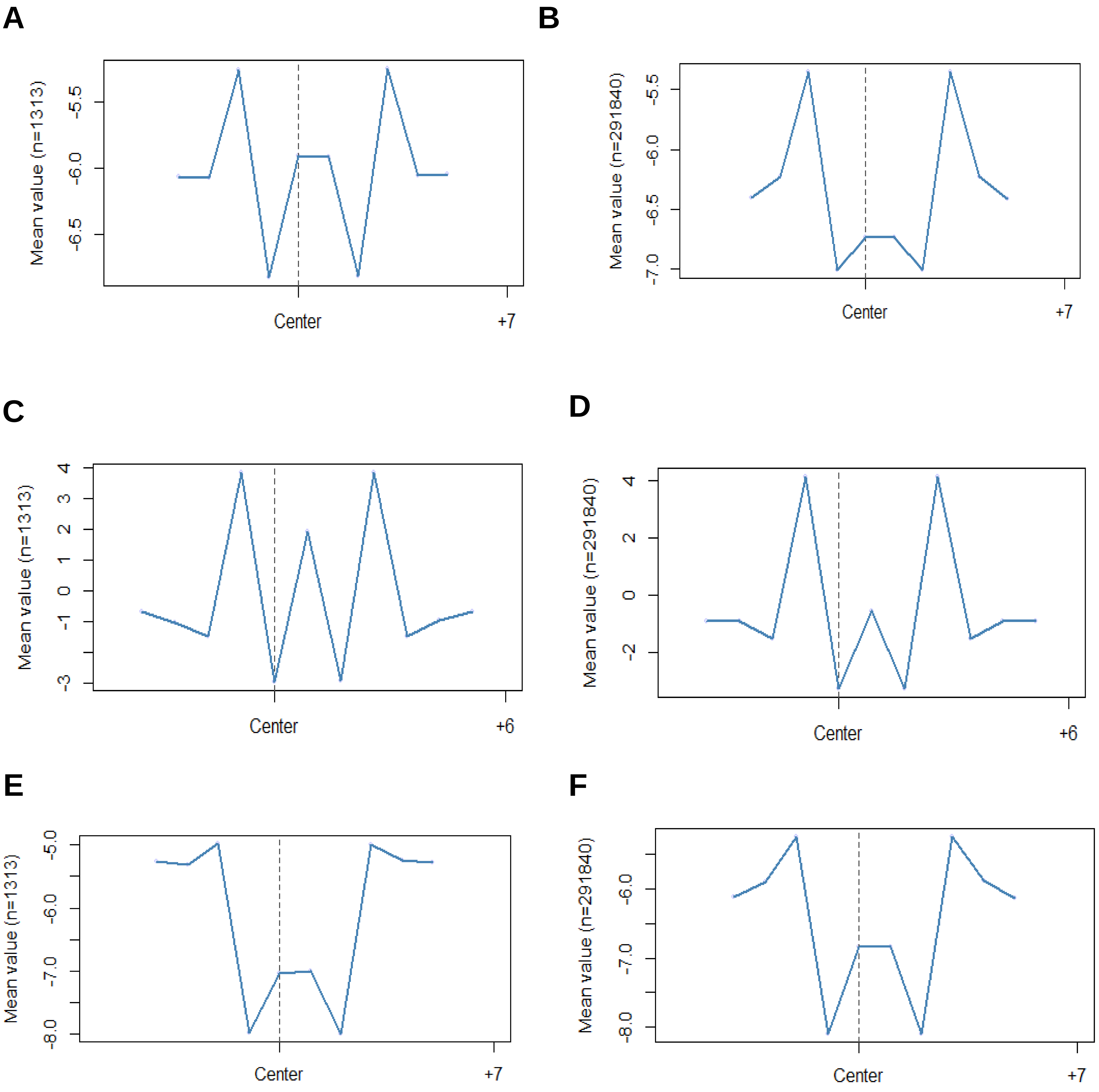


Supplementary Figure 5: **Heart DNA Shape features.** Electrostatic Potential (EP) at **(A)** BMAL1 bound (**B)** BMAL1 unbound E-boxes. Roll at **(C)** BMAL1 bound **(D)** BMAL1 unbound E-boxes. Propeller Twist (ProT) at **(E)** BMAL1 bound **(F)** BMAL1 unbound E-boxes. The differences in the bound and unbound shape features contribute to the model accurately predicting BMAL1 binding to the E-box motifs. The mean values represent the number of E-box motifs used in generating the shape figure.

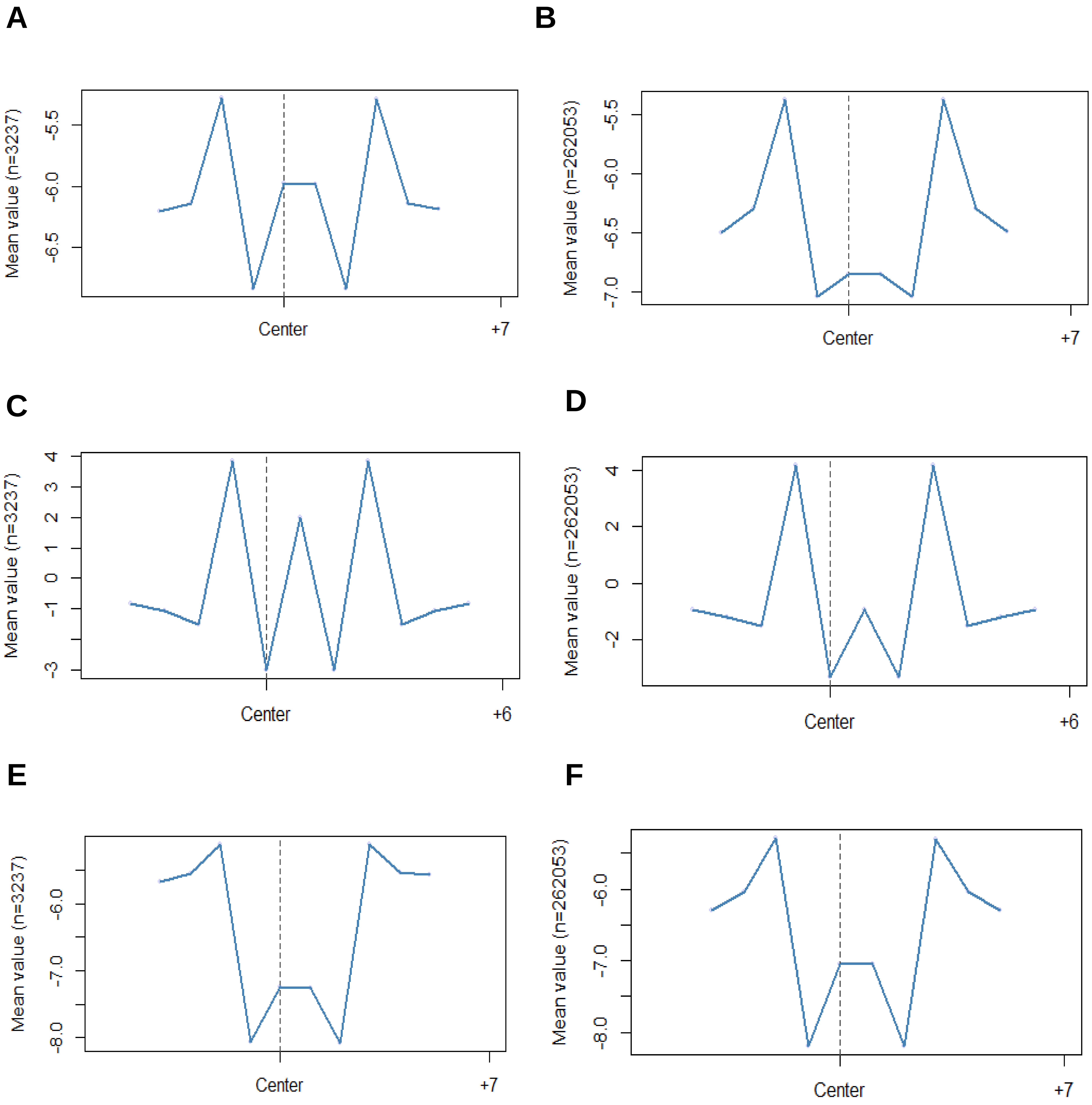


• Supplementary Figure 6**: Kidney DNA Shape features.** Electrostatic Potential (EP) at **(A)** BMAL1 bound **(B)** BMAL1 unbound E-boxes. Roll at **(C)** BMAL1 bound **(D)** BMAL1 unbound E-boxes. Propeller Twist (ProT) at **(E)** BMAL1 bound **(F)** BMAL1 unbound E-boxes. The differences in the bound and unbound shape features contribute to the model accurately predicting BMAL1 binding to the E-box motifs. The mean values represent the number of E-box motifs used in generating the shape figure.

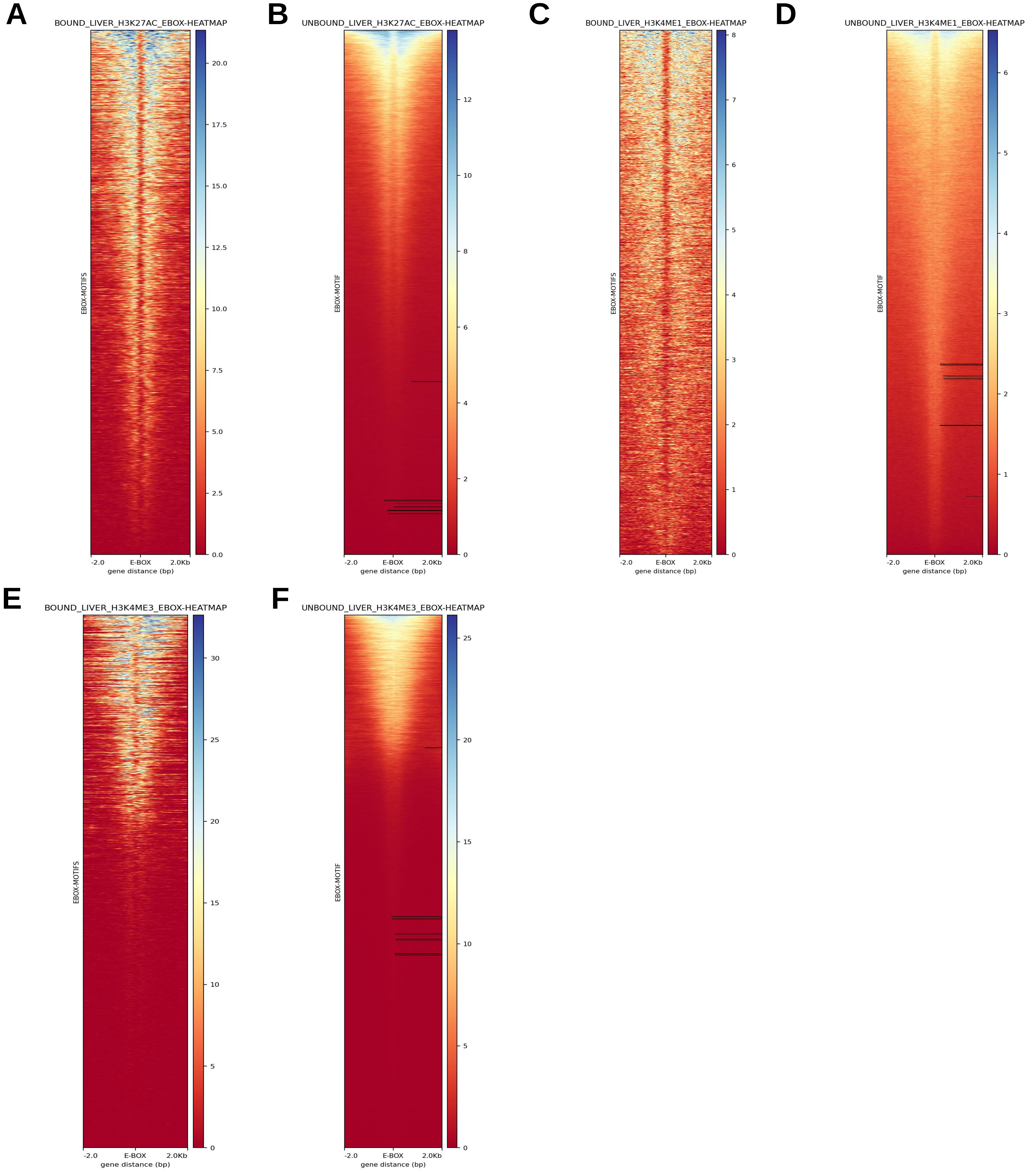


Supplementary Figure 7: **Liver histone modification profiles.** H3K27ac at **(A)** BMAL1 bound **(B)** BMAL1 unbound E-boxes. H3K4ME1 at **(C)** BMAL1 bound **(D)** BMAL1 unbound E-boxes. H3K4ME3 at **(E)** BMAL1 bound **(F)** BMAL1 unbound E-boxes. The differences in the bound and unbound profile shapes contribute to the model accurately predicting BMAL1 binding to the E-box motifs.

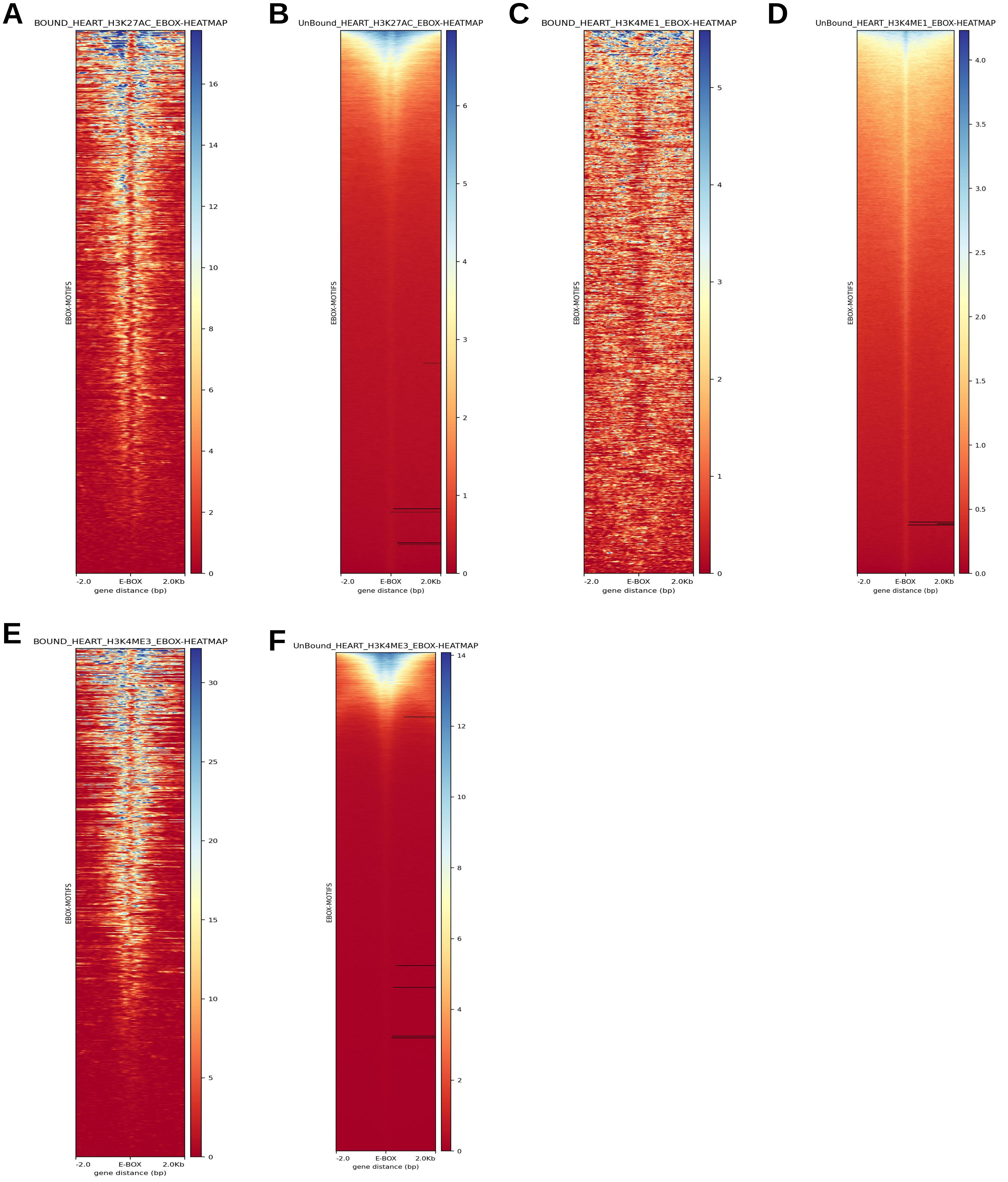


Supplementary Figure 7: **Heart histone modification profiles.** H3K27ac at **(A)** BMAL1 bound **(B)** BMAL1 unbound E-boxes. H3K4ME1 at **(C)** BMAL1 bound **(D)** BMAL1 unbound E-boxes. H3K4ME3 at **(E)** BMAL1 bound **(F)** BMAL1 unbound E-boxes. The differences in the bound and unbound profile shapes contribute to the model accurately predicting BMAL1 binding to the E-box motifs.

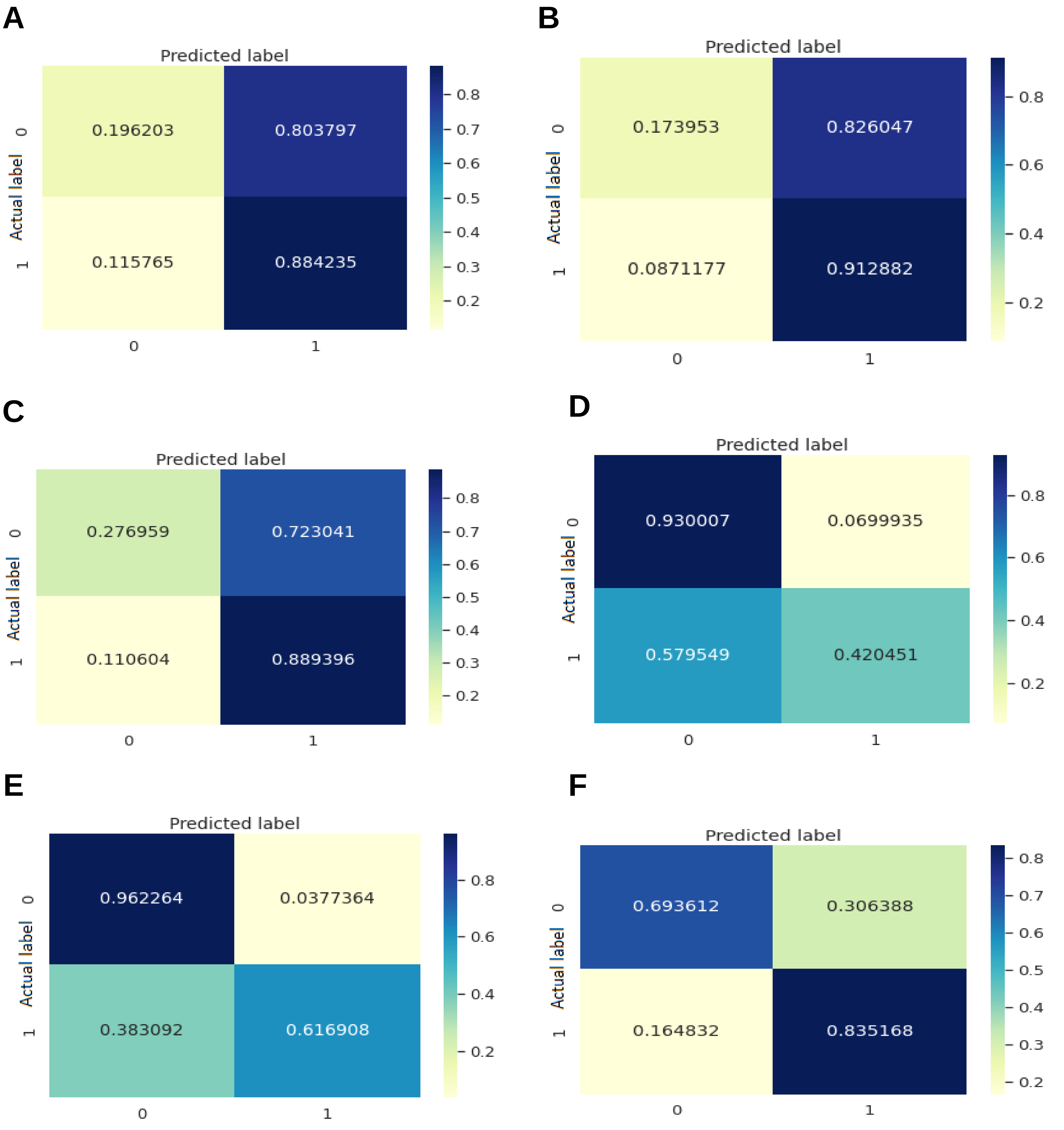


• Supplementary Figure 9: **Confusion matrix for cross tissue models. (A)** Liver_Heart **(B)** Liver_Kidney **(C)** Heart_Liver **(D)** Heart_Kidney **(E)** Kidney_Heart **(F)** Kidney_Liver (Liver-Heart means the model was trained on the all the liver dataset and used to predict binding in the heart dataset.)
